# The relationship between longevity and diet is genotype dependent and sensitive to desiccation

**DOI:** 10.1101/2020.01.16.908996

**Authors:** Andrew W McCracken, Eleanor Buckle, Mirre J. P. Simons

**Affiliations:** Department of Animal and Plant Sciences & Bateson Centre, The University of Sheffield, Sheffield S10 2TN, UK

**Keywords:** Dietary restriction, Drosophila melanogaster, reaction norm, overfeeding, starvation, desiccation

## Abstract

Dietary restriction (DR) is a key focus in ageing research. Specific conditions and genotypes were recently found to negate lifespan extension by DR, questioning its universal relevance. However, the conceptual framework of dietary reaction norms explains why DR’s effects might not be apparent in some situations. We tested comprehensively the importance of dietary reaction norms by measuring longevity and fecundity on five diets in five genotypes, with and without water supplementation in the fly (N>25,000). We found substantial genetic variation in the reaction norm between diet and lifespan. Environments supplemented with water rescued putative desiccation stress but only at the richest diets. Fecundity declined at these richest diets, but was unaffected by water and is thus most likely caused by nutritional toxicity. Our results demonstrate empirically that any conclusion on the absence of DR is only justified when a range of diets is considered in a reaction norm framework.

## Introduction

Dietary restriction (DR), the limitation of food intake but avoiding malnutrition, extends lifespan. The generality of the DR response has been questioned, however, by reports that DR does not extend lifespan under certain experimental conditions (Austad, 2012; Dick et al., 2011; Ja et al., 2009 [but see Piper et al., 2010]) or in a considerable proportion of the genotypes tested (Dick et al., 2011; Liao et al., 2010; Mitchell et al., 2016; Rikke et al., 2010; Swindell, 2012; Wilson et al., 2018 preprint). These conclusions are routinely based upon experiments using two diets (dietary dyad) alone, whereas it is recognised that a change in the continuous relationship between diet and lifespan (reaction norm) can obscure lifespan extension by DR (Flatt, 2014; Tatar, 2011). The bell-shaped nature of the dietary reaction norm dictates that one particular diet concentration, in one genotype or environment, will result in the longest lifespan; lower or higher diet concentrations will induce a shortened lifespan due to malnutrition or overfeeding, respectively. Where a particular dietary dyad falls on this reaction norm will determine the magnitude of the DR effect and can even lead to the erroneous conclusion that DR shortens lifespan (Fig.1).

**Fig.1.**
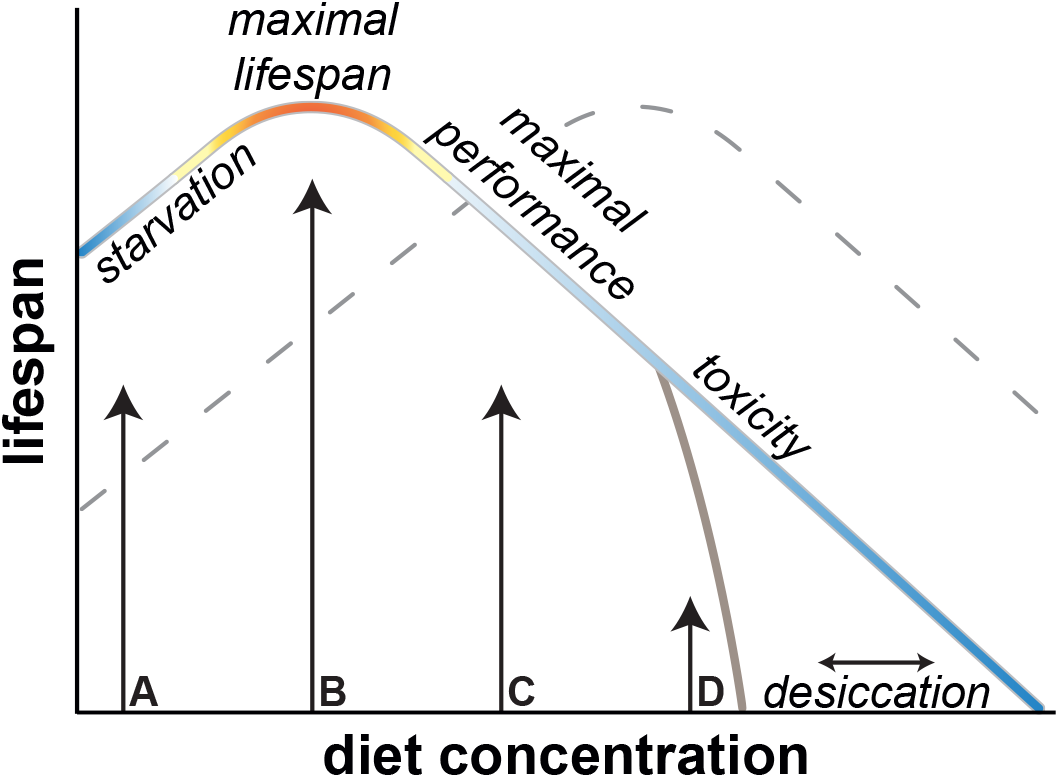
Schematic of multiple thresholds in the lifespan reaction norm to diet. Diet concentration has a bell-shaped relationship with lifespan, ranging from malnutrition (A), DR (B), maximal performance - or highest Darwinian fitness - at a relatively rich diet (C), to overfeeding, leading to nutritional toxicity (D). As a detailed reaction norm is rarely known, a dietary dyad although often used can lead to misleading conclusions. A dietary dyad (A & C) can show no response at all due to the symmetry in the shape of the reaction norm. Furthermore, genetic or environmental effects can alter the shape or shift the reaction norm (dashed line), or lead to effects at only specific parts of the reaction norm (solid gray, e.g. desiccation). For example, diets B & C result in a DR response on the focal curve, but malnutrition on the dashed curve.

Few studies have examined dietary reaction norms in more detail by titrating the supply of protein or calories across multiple genotypes or environments, and none have tested both genetic and environmental effects on dietary reaction norms simultaneously. Of these studies, a proportion employed transgenic or lab strains (Clancy et al., 2002; Grandison et al., 2009; Min et al., 2008; Skorupa et al., 2008; Tatar, 2011; Wang et al., 2009) and demonstrated varying degrees of genetic variance in the plastic response to diet. Across these studies, shifts in dietary reaction norms on the x- or y-plane are more apparent than changes in the overall shape of the relationship between diet and longevity (Flatt, 2014; Tatar, 2011). Whether genetic variation in transgenic and lab strain experiments is representative of standing genetic variation of natural populations is, however, unclear. A naturalistic appreciation of the genetic variation of the DR response becomes particularly important when null responses are interpreted to question the universal properties of DR important in translating its benefits to our own species. One previous study did measure detailed reaction norms using wild-derived outbred populations and found a degree of genetic variance for the relationship between diet and lifespan (Metaxakis and Partridge, 2013). However, the estimate of genetic variance of a population level trait, such as lifespan, when estimated from between outbred stains (Whitlock and Fowler, 1999) will be affected by mortality heterogeneity (Chen et al., 2013), which can bias the estimated level of genetic variance upwards or downwards.

When specific environmental effects interact or interfere with the DR reaction norm, the use of dietary dyads - or the neglect of environmental confounds - could similarly lead to misleading conclusions. For flies specifically, water supplementation has been suggested to diminish the effect of DR on lifespan (Dick et al., 2011; Ja et al., 2009). The conclusion that water completely explains DR has been discredited (Piper et al., 2010), but flies nonetheless consider water a nutrient and consume 1-2μl per day, with higher consumption at higher dietary yeast (Fanson et al., 2012) and sugar concentrations (van Dam et al., 2020). Hence, erroneous conclusions could be drawn from diet responses if desiccation presents a genotype- or diet-specific hazard.

Here, we present DR reaction norms for fecundity and longevity across five genotypes in female flies (*Drosophila melanogaster*) with and without water supplementation using high sample sizes. We show empirically across five wild-derived, inbred lines that there are strong genetic and environmental elements to dietary reaction norms, and therefore the thorough appreciation of reaction norms is critical when interpreting diet effects across genotypes and environments.

## Materials and Methods

### Fly husbandry, experimental protocol and dietary regimes

For lifespan experiments adults were provided with either 0.5%, 2%, 5%, 8% or 14% autolysed yeast media. All other media components (13% table sugar, 6% cornmeal, 1% agar and 0.225% [w/v] nipagin)l remained the same, given the dietary protein axis is the main lifespan determinant in flies (Jensen et al., 2015; Lee et al., 2008). Note, cornmeal concentration was halved in 14% yeast media to allow dispensing of the media. Purpose-built demography cages included two openings, one for the supplementation of food, and one for water-agar (2% agar) or empty vial. Cages contained between 70-125 females each (mode of ~ 100 females), with 5 cages per treatment, per genotype (N = 50 cages per genotype). For one genotype, DGRP-195, sample size was even higher: an additional two cages of water-supplemented, and control cages at 2% media. All experimental flies were mated on 8% media for 48 hours, and kept in cages on 8% media until age 3-4 days, when experimental dietary treatments started.

To establish dietary reaction responses, flies were exposed to continuous diets with the addition, or absence of water-agar supplementation. To test the effect of water supplementation on longevity, we provided an additional vial of water-agar (‘water supplementation’), or an empty vial (‘control’), to each cage. Separation of food and water sources allowed flies to choose their source of nourishment, and eliminated the need for hydration to be coupled with caloric intake. Dietary treatments were balanced for age, and date of eclosion. All flies presented were grown within one batch. The experiment was carried out on a small collection of DGRP lines (Mackay et al., 2012); DGRP-195; 217; 239; 362; 853).

### Fecundity

Feeding vials were imaged and analysed using image analysis software QuantiFly (Waithe et al., 2015) to determine the relative quantity of egg laying. Vials were removed, during normal scoring periods, from all cages containing eggs from flies aged 11 or 12.

### Data analysis

Mixed Cox-proportional hazard models were used that included ‘cage’ as random term to correct for uncertainty of pseudo-replicated effects within demography cages (Ripatti and Palmgren, 2000; Therneau et al., 2003). Additional specific tests of coefficients are provided that combine the single and interaction term (in a z-test, using the maximum s.e. of the factors compared) to test how survival was changing in water-treated flies, compared to respective control treatments. Note, formal tests for proportionality of hazards are not available for mixed effects Cox regressions. For survival data comparisons, we report the full model, and models fitted within each genotype separately (see supplement). By splitting the analysis between genotypes, bias introduced by deviations in proportionality of hazards between genotypes is avoided. Qualitative conclusions remain similar, irrespective of how these models are fitted. Coefficients are reported as logged hazards with significance based on z-tests. Right-censoring was included, and dietary treatments were considered categorical factors.

Egg laying was analysed using linear models of log-transformed fecundity count data. Flies only differed by one day in age, and age was equally distributed across treatment and measured in a balanced design. BIC with backward elimination of terms was used for model comparisons and selection, and resulted in a model that only contained the interaction between genotype and diet. Water was added to our models to directly test for any effect on fecundity, but this proved negligible (Table S13,14).

For hazard ratio figures, ratios are plotted as coefficients derived from within-line Cox mixed-effect models, with error bars representing 95% confidence intervals.

## Results and Discussion

The genotypes tested showed a classical bell-shaped response to diet. Longer lifespans were observed at intermediate dietary yeast concentrations, consistent with DR (Fig.2A,S4; Table S1-6; P < 0.001). All genotypes also exhibited a reduction in survival at very lowest yeast concentrations (starvation), and at the very highest (maximal performance or nutritional toxicity). We detected considerable genetic variation in the response to diet (χ2=162, df=16, P< 0.001) with the diet of maximum longevity differing between genotypes.

**Fig.2.**
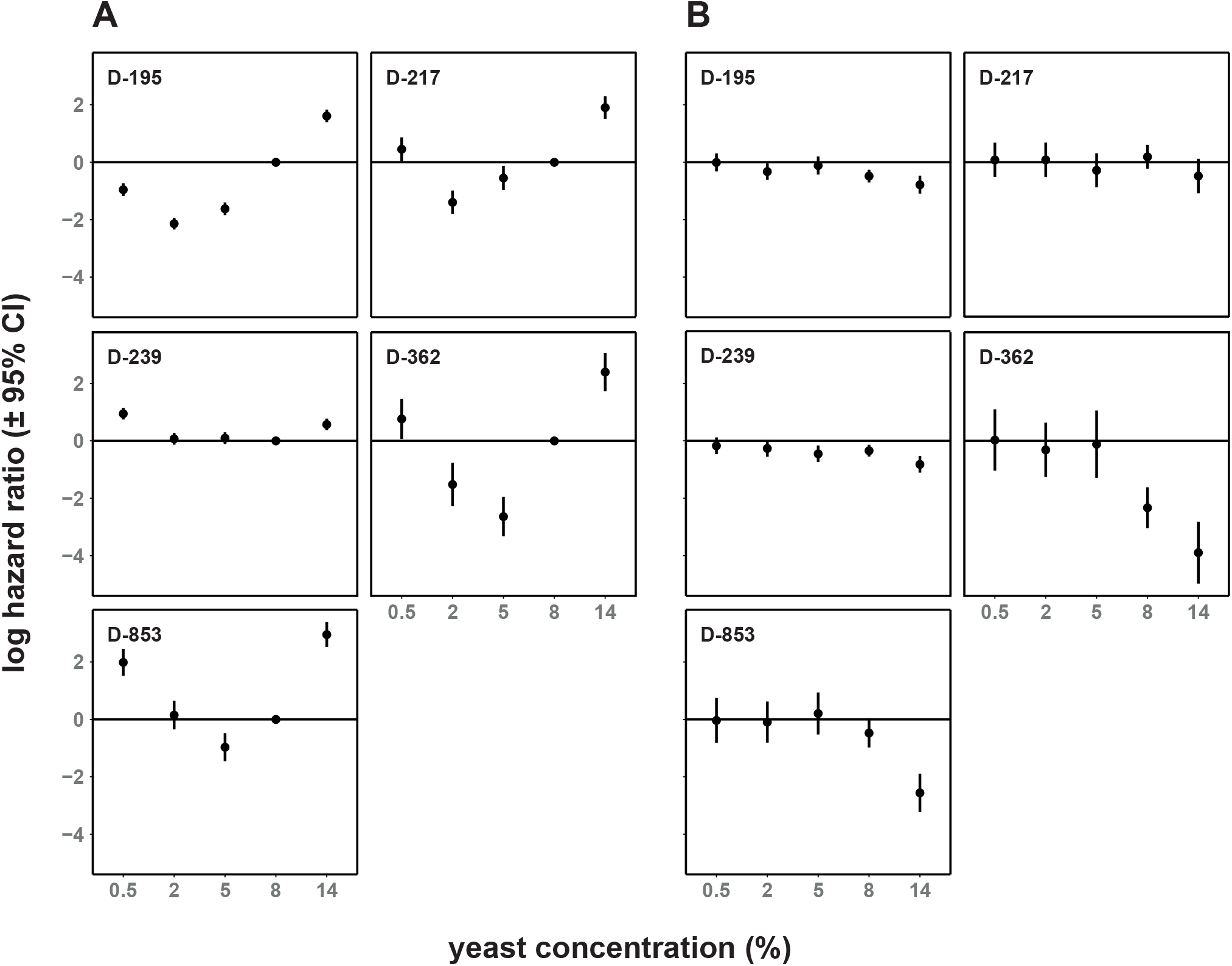
Log hazard ratios of diet and water supplementation in a panel of DGRP genotypes. **A -** Dietary reaction norms vary in a genotype-specific manner. **B -** Water-supplementation, relative to control treatment, rescues desiccation in a diet and genotype-dependent manner. Hazard ratios represent risk to die, therefore higher values indicate shorter lifespans and are relative. Ratios are plotted as coefficients derived from within-line Cox mixed-effect models, with error bars representing 95% confidence intervals. For **A**, 8% yeast treatment was treated as a reference and as such, no CIs are available. Rates here are relative to 8% yeast diet, and lines represent this standard. N = 25,519 females total; 4,800-5,282 per genotype. For **B**, hazard rates are relative to the corresponding control for each diet. Horizontal lines represent a water effect size of 0. N = 12,737 females total; 2,396-2629 per genotype.

To test the effect of desiccation, we compared longevity under control conditions to water-supplemented. Supplemental water reduced mortality particularly at higher yeast concentrations, and we found genetic variance for this environmental effect (χ2=160, df=16, P<0.001; Fig.2B; Table S2-6). At the highest yeast concentrations, this amounted to a one and a half-to fifty-fold reduction in hazard rate. Given this, particular caution should be afforded when considering the effect of desiccation, especially in organisms without ad libitum access to water and when fed a concentrated diet. To assess statistically whether water supplementation abolished DR-induced life extension (Ja et al., 2009; Piper et al., 2010) we ran our statistical models within the water treatment only, but found no evidence for this suggestion (Fig.S1, Table S7-12). Given that water supplementation ameliorated, but did not eliminate, elevated mortality under the highest yeast concentrations, we conclude desiccation can play an experimentally confounding role in DR, but is not causal.

DR is known to reduce reproductive output and is interpreted as a response to decreased energy availability (Moatt et al., 2016). The effect of overfeeding on reproduction, although appreciated in humans (Broughton and Moley, 2017), has received little attention (McCracken et al., 2020). These two responses were evident in egg laying: an increase with yeast concentration, and a stabilisation, or decline at the highest yeast concentrations (Fig.S2,3; Table S13,14). As with mortality, genetic lines also differed in the reproductive response to diet (F=6.3, df=16, P< 0.001). Reduced egg laying together with a reduction in survival, lowered predicted lifetime reproductive at the richest diet (Fig.S3). Egg laying was not affected by water supplementation (Fig.S2; Table S13,14; P>0.15. Notably, even when water rescued mortality caused by desiccation at the high yeast concentrations, egg laying was unaffected (Fig.2,S2). Given this, we infer the decline in reproductive output at the highest yeast concentration was not due to desiccation stress, but nutritional toxicity. By contrast, the rescue of mortality at high yeast concentrations by water supplementation is therefore likely to be separate and driven by desiccation.

In conclusion, we observe significant genetic, and environmentally induced, variation in the lifespan and fecundity responses to diet. These data now directly demonstrate that specific care is needed when interpreting effects of DR across genotypes, experimental conditions or environments. We acknowledge that carrying out full reaction norms in all DR experiments would be laborious. Therefore, we suggest selecting dietary dyads that differ only minimally when genetic variance in DR is the object of study. Such a strategy reduces the chance that tested diets diverge considerably from maximal lifespans, leading to starvation or nutritional toxicity (Fig.1). Furthermore, we suggest when environmental conditions, such as water (Ja et al., 2009), sex (Regan et al., 2016) and microbiome (Wong et al., 2014) are presumed to negate the DR response, that a post-hoc reaction norm is performed. Similar considerations hold for mechanistic research. Should, for example, a genetic manipulation remove the DR response, only a full dietary reaction norm can demonstrate how such an effect arises: by either a shift in, or compression of, the reaction norm (Flatt, 2014; Tatar, 2011). The importance of reaction norms when studying DR has been stressed before, but this is the first high sample size data across multiple wild-type inbred genotypes and diets, including an environmental confound, that demonstrates this empirically.

## Acknowledgements

We thank the Deplancke laboratory for supplying DGRP lines and members of Simons’ lab for their valuable assistance.

## Author Contributions

AWM and MJPS designed and interpreted the experiments. AWM drafted the first version of the manuscript. AWM and MJPS revised the manuscript. AWM led the data acquisition with help from EB. MJPS supervised the project.

## Supplementary information

### Figures

**Fig. S1.**
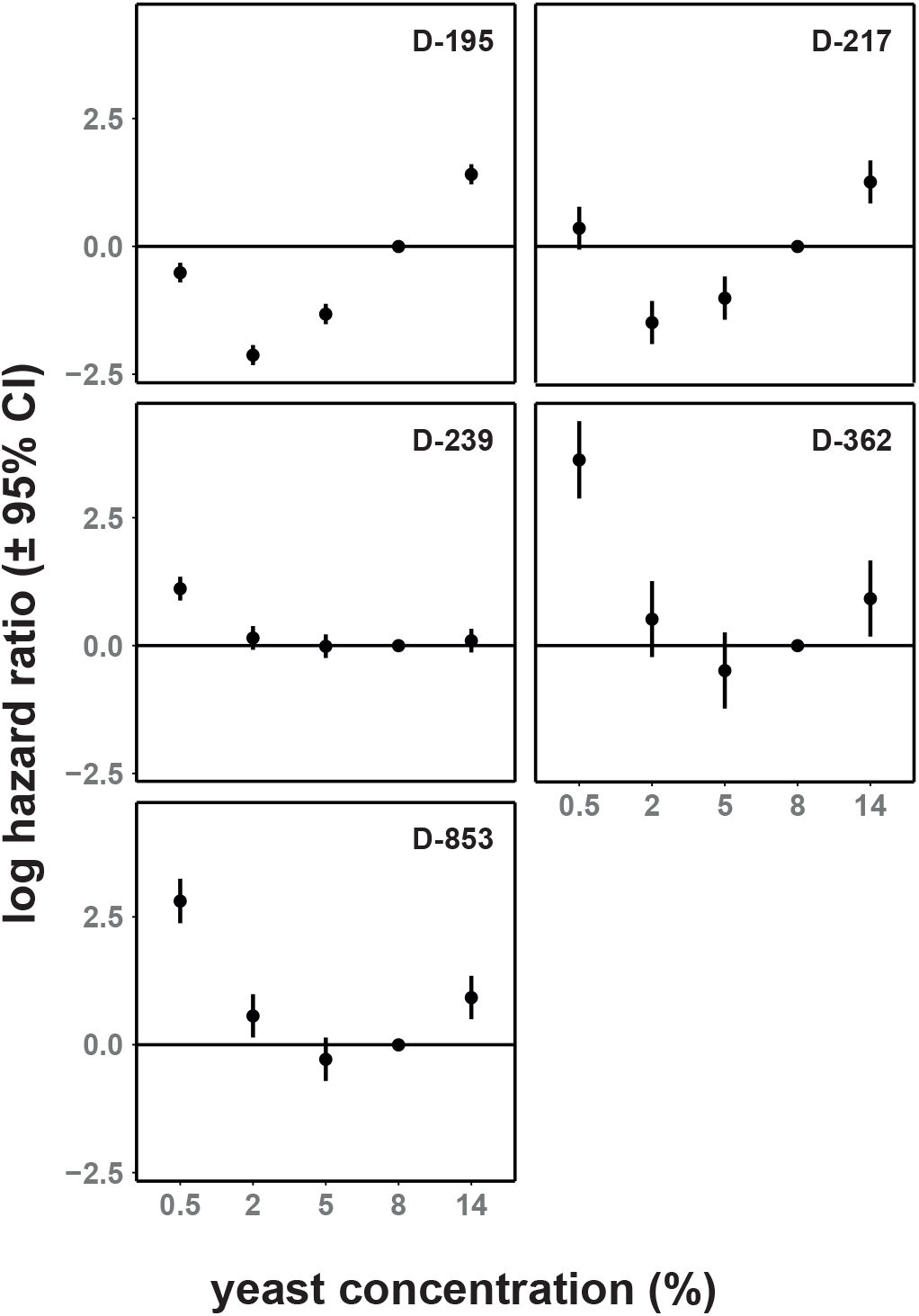
Log hazard ratios of diet within water-treated cages in a panel of DGRP genotypes. Reaction norms to diet still differ in water-treated circumstances. Hazard ratios represent the inverse of typical survival reaction norms to diet. 8% yeast treatment was treated as a reference and as such, no CIs are available. Rates here are relative to 8% yeast diet, and lines represent this standard. N = 12,737 females total; 2,396-2629 per genotype. Hazard ratios have the benefit over median lifespan in that they are directly related to the appropriate statistics used for time-to-event data. In addition, they are directly comparable in a quantitative fashion across genotypes of different lifespans, as they express a relative risk.

**Fig. S2.**
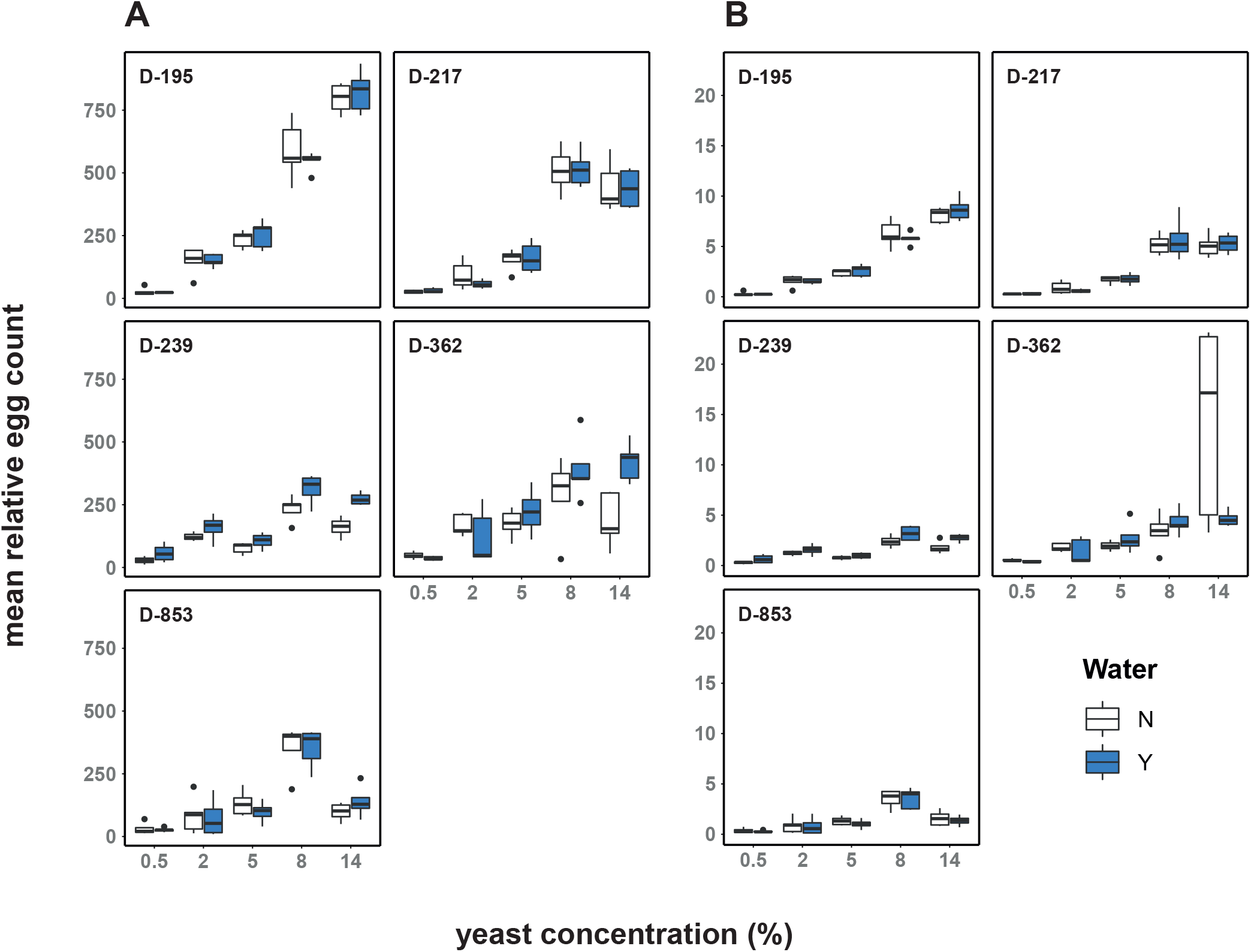
Fecundity analysis of panel under all conditions. Fecundity has a positive relationship with dietary yeast concentration, except at the highest yeast concentration assayed (14%) for most genotypes. **A** - raw egg counts. **B** - mortality-corrected counts. Counts generated using QuantiFly software. Counts are relative, but directly comparable. Flies assayed at age 11-12 days, with boxplots aggregating totals (median, with the box depicting a quartile each way, and whiskers showing the range; outliers plotted as dots). Each cage was assayed on 1 scoring day at this age. Mortality corrected counts (**B**) generated by dividing raw counts, by N flies remaining in cage at the time of assaying. N = 25,519 females total; 4,800-5,282 per genotype. Note that DGRP-362 experienced significant mortality at this age under 14% yeast dietary treatment. This is the cause of the discrepancy in significance between raw, and age-adjusted fecundity counts. Note, egg-laying was not assessed throughout life and in natural circumstances lifespan of the fly is truncated by extrinsic factors, e.g. predation. We nonetheless, tentatively conclude that the enhanced mortality and reduced egg laying on very rich diets is caused by nutritional toxicity.

**Fig. S3.**
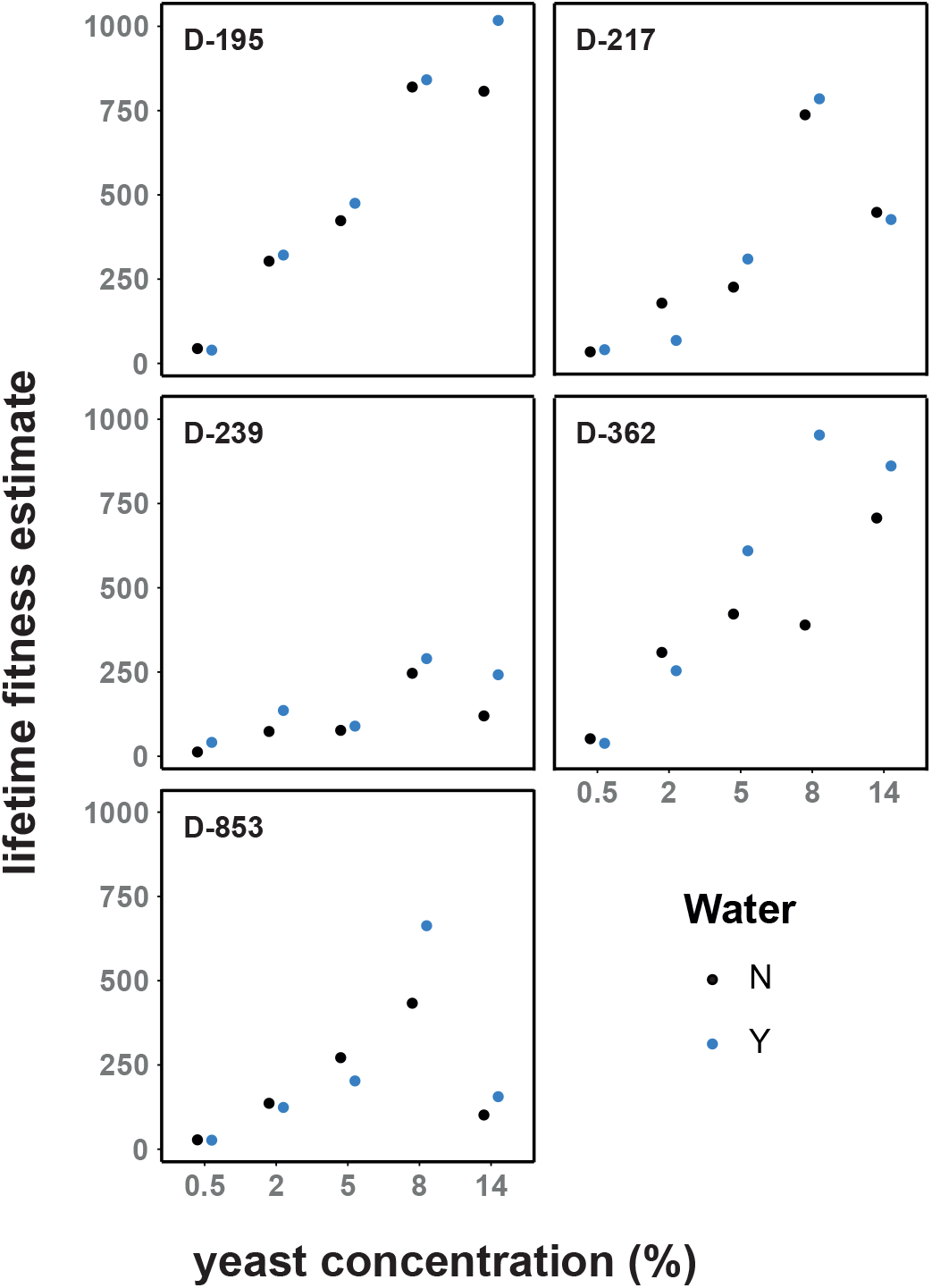
Lifetime reproductive fitness estimates of panel under all conditions. Lifetime fitness has a positive relationship with dietary yeast concentration, except at the highest yeast concentration assayed (14%) for most genotypes. Mortality-adjusted egg counts from Fig. S2 were multiplied by the area under the relevant survival curve (restricted mean) to generate lifetime estimates. N = 25,519 females total; 4,800-5,282 per genotype.

**Fig. S4.**
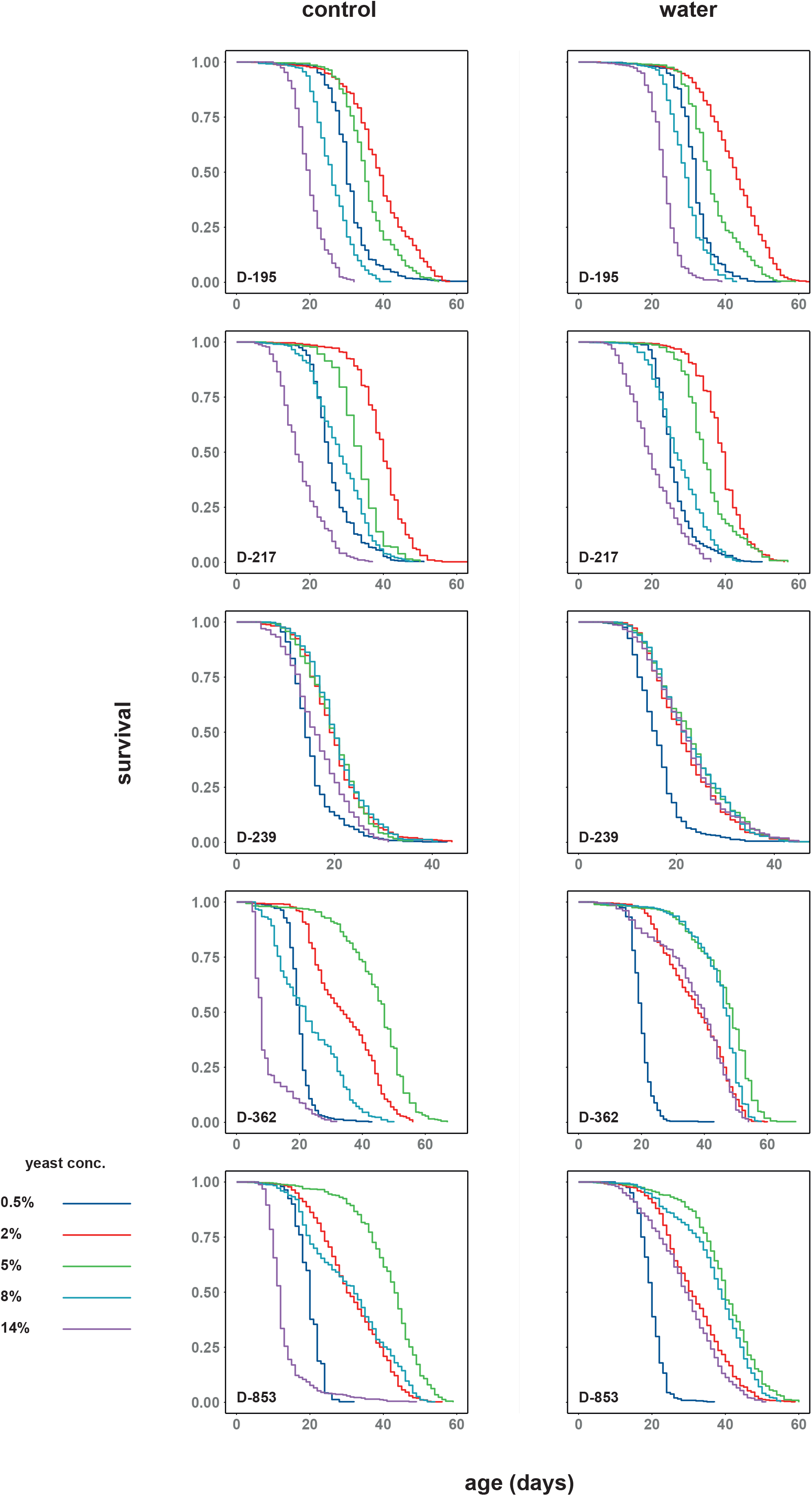
Survival curves of panel in response to diet. Dietary reaction norms vary in a genotype-specific manner. Survival curves are separated by genotype, and water-supplementation status. N = 25,519 females total; 4,800-5,282 per genotype.

### Tables

**Table S1.** Effect of diet and water supplementation on mortality across 5 DGRP lines (DGRP-195 is reference).

| coefficient | Full Model |  |  |  |
| --- | --- | --- | --- | --- |
|  | estimate | exp | s.e. | p |
| water | -0.390 | 0.677 | 0.228 | 0.086 |
| 0.5% yeast | -0.796 | 0.451 | 0.228 | <0.001 |
| 2% yeast | -1.856 | 0.156 | 0.206 | <0.001 |
| 5% yeast | -1.296 | 0.274 | 0.225 | <0.001 |
| 14% yeast | 1.043 | 2.837 | 0.226 | <0.001 |
| 217 | -0.444 | 0.641 | 0.226 | 0.05 |
| 239 | 0.763 | 2.145 | 0.227 | 0.001 |
| 362 | 0.104 | 1.110 | 0.227 | 0.645 |
| 853 | -1.090 | 0.336 | 0.223 | <0.001 |
| 0.5% yeast * water | 0.470 | 1.601 | 0.322 | 0.144 |
| 2% yeast * water | 0.030 | 1.031 | 0.300 | 0.92 |
| 5% yeast * water | 0.277 | 1.319 | 0.322 | 0.39 |
| 14% yeast * water | -0.156 | 0.856 | 0.322 | 0.629 |
| 217 * 0.5% yeast | 1.195 | 3.302 | 0.320 | <0.001 |
| 217 * 2% yeast | 0.632 | 1.881 | 0.310 | 0.042 |
| 217 * 5% yeast | 0.786 | 2.195 | 0.321 | 0.014 |
| 217 * 14% yeast | 0.643 | 1.903 | 0.322 | 0.046 |
| 239 * 0.5% yeast | 1.793 | 6.008 | 0.321 | <0.001 |
| 239 * 2% yeast | 1.887 | 6.599 | 0.308 | <0.001 |
| 239 * 5% yeast | 1.429 | 4.174 | 0.321 | <0.001 |
| 239 * 14% yeast | -0.382 | 0.683 | 0.324 | 0.238 |
| 362 * 0.5% yeast | 1.754 | 5.775 | 0.319 | <0.001 |
| 362 * 2% yeast | 0.262 | 1.300 | 0.311 | 0.399 |
| 362 * 5% yeast | -1.322 | 0.267 | 0.315 | <0.001 |
| 362 * 14% yeast | 1.880 | 6.551 | 0.316 | <0.001 |
| 853 * 0.5% yeast | 3.087 | 21.901 | 0.318 | <0.001 |
| 853 * 2% yeast | 2.029 | 7.604 | 0.307 | <0.001 |
| 853 * 5% yeast | 0.287 | 1.332 | 0.320 | 0.37 |
| 853 * 14% yeast | 2.100 | 8.168 | 0.320 | <0.001 |
| 217 * water | 0.551 | 1.736 | 0.322 | 0.087 |
| 239 * water | -0.101 | 0.904 | 0.322 | 0.754 |
| 362 * water | -2.015 | 0.133 | 0.319 | <0.001 |
| 853 * water | -0.101 | 0.904 | 0.323 | 0.753 |
| 217 * 0.5% yeast * water | -0.568 | 0.567 | 0.455 | 0.212 |
| 217 * 2% yeast * water | -0.114 | 0.893 | 0.441 | 0.797 |
| 217 * 5% yeast * water | -0.702 | 0.496 | 0.456 | 0.123 |
| 217 * 14% yeast * water | -0.448 | 0.639 | 0.456 | 0.327 |
| 239 * 0.5% yeast * water | -0.145 | 0.865 | 0.457 | 0.751 |
| 239 * 2% yeast * water | 0.126 | 1.134 | 0.440 | 0.775 |
| 239 * 5% yeast * water | -0.427 | 0.652 | 0.455 | 0.348 |
| 239 * 14% yeast * water | -0.383 | 0.682 | 0.457 | 0.402 |
| 362 * 0.5% yeast * water | 1.990 | 7.319 | 0.456 | <0.001 |
| 362 * 2% yeast * water | 2.068 | 7.912 | 0.439 | <0.001 |
| 362 * 5% yeast * water | 2.007 | 7.444 | 0.457 | <0.001 |
| 362 * 14% yeast * water | -1.942 | 0.143 | 0.451 | <0.001 |
| 853 * 0.5% yeast * water | -0.027 | 0.974 | 0.458 | 0.954 |
| 853 * 2% yeast * water | 0.351 | 1.420 | 0.441 | 0.427 |
| 853 * 5% yeast * water | 0.433 | 1.541 | 0.457 | 0.344 |
| 853 * 14% yeast * water | -2.056 | 0.128 | 0.455 | <0.001 |

**Table S2.** Effect of diet and water supplementation on mortality within DGRP-195.

| coefficient | Estimates from individual model |  |  |  | Effect of water, compared to no water |  |  |
| --- | --- | --- | --- | --- | --- | --- | --- |
|  | estimate | exp | s.e. | p | estimate | exp | p |
| water supplementation | -0.481 | 0.618 | 0.113 | <0.001 |  |  |  |
| 0.5% yeast | -0.954 | 0.385 | 0.112 | <0.001 |  |  |  |
| 2% yeast | -2.139 | 0.118 | 0.103 | <0.001 |  |  |  |
| 5% yeast | -1.620 | 0.198 | 0.112 | <0.001 |  |  |  |
| 14% yeast | 1.612 | 5.012 | 0.112 | <0.001 |  |  |  |
| 0.5% yeast * water | 0.475 | 1.608 | 0.158 | 0.003 | -0.006 | 0.994 | 0.971 |
| 2% yeast * water | 0.156 | 1.169 | 0.147 | 0.289 | -0.324 | 0.723 | 0.028 |
| 5% yeast * water | 0.369 | 1.446 | 0.160 | 0.021 | -0.112 | 0.894 | 0.483 |
| 14% yeast * water | -0.301 | 0.740 | 0.159 | 0.059 | -0.781 | 0.458 | <0.001 |

**Table S3.** Effect of diet and water supplementation on mortality within DGRP-217.

| coefficient | Estimates from individual model |  |  |  | Effect of water, compared to no water |  |  |
| --- | --- | --- | --- | --- | --- | --- | --- |
|  | estimate | exp | s.e. | p | estimate | exp | p |
| water supplementation | 0.190 | 1.209 | 0.213 | 0.373 |  |  |  |
| 0.5% yeast | 0.454 | 1.574 | 0.211 | 0.032 |  |  |  |
| 2% yeast | -1.395 | 0.248 | 0.208 | <0.001 |  |  |  |
| 5% yeast | -0.549 | 0.578 | 0.212 | 0.01 |  |  |  |
| 14% yeast | 1.905 | 6.716 | 0.200 | <0.001 |  |  |  |
| 0.5% yeast * water | -0.108 | 0.898 | 0.306 | 0.724 | 0.082 | 1.08 | 0.789 |
| 2% yeast * water | -0.105 | 0.901 | 0.306 | 0.732 | 0.085 | 1.08 | 0.781 |
| 5% yeast * water | -0.470 | 0.625 | 0.300 | 0.117 | -0.280 | 0.75 | 0.350 |
| 14% yeast * water | -0.668 | 0.513 | 0.306 | 0.029 | -0.479 | 0.62 | 0.118 |

**Table S4.** Effect of diet and water supplementation on mortality within DGRP-239.

| coefficient | Estimates from individual model |  |  |  | Effect of water, compared to no water |  |  |
| --- | --- | --- | --- | --- | --- | --- | --- |
|  | estimate | exp | s.e. | p | estimate | exp | p |
| water supplementation | -0.343 | 0.710 | 0.105 | 0.001 |  |  |  |
| 0.5% yeast | 0.946 | 2.575 | 0.104 | <0.001 |  |  |  |
| 2% yeast | 0.070 | 1.072 | 0.104 | 0.502 |  |  |  |
| 5% yeast | 0.094 | 1.098 | 0.103 | 0.363 |  |  |  |
| 14% yeast | 0.572 | 1.771 | 0.103 | <0.001 |  |  |  |
| 0.5% yeast * water | 0.173 | 1.188 | 0.148 | 0.243 | -0.170 | 0.843 | 0.249 |
| 2% yeast * water | 0.079 | 1.082 | 0.148 | 0.593 | -0.264 | 0.768 | 0.075 |
| 5% yeast * water | -0.108 | 0.898 | 0.147 | 0.464 | -0.451 | 0.637 | 0.002 |
| 14% yeast * water | -0.476 | 0.621 | 0.147 | 0.001 | -0.819 | 0.441 | <0.001 |

**Table S5.**
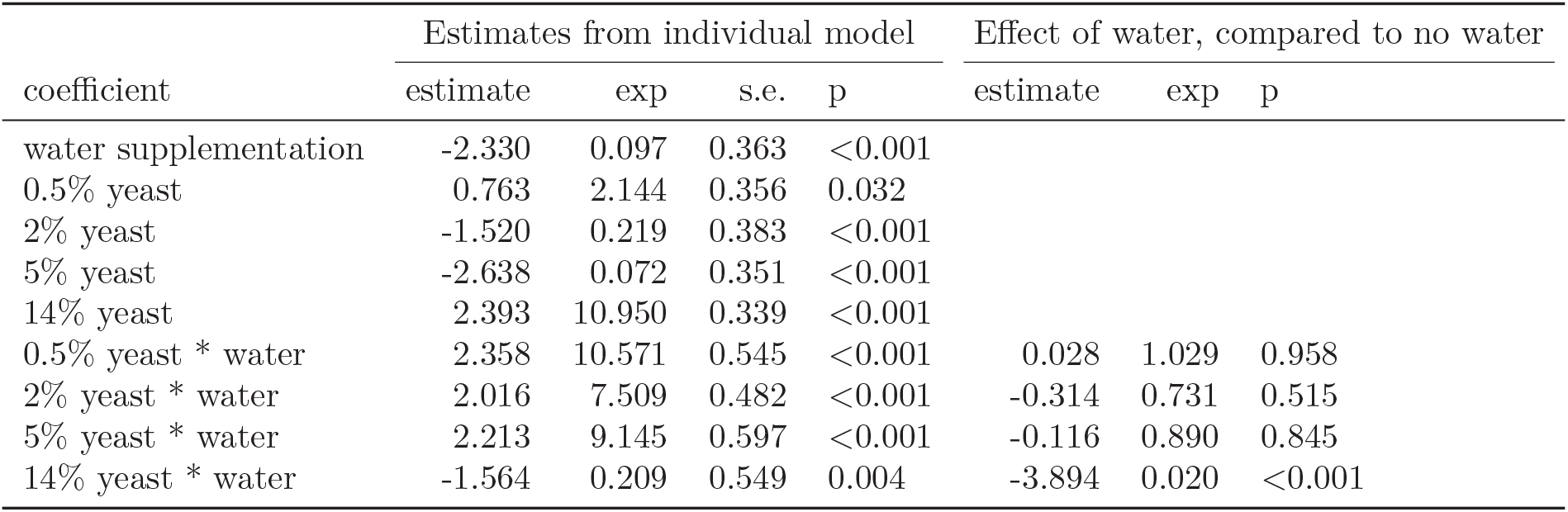
Effect of diet and water supplementation on mortality within DGRP-362.

| coefficient | Estimates from individual model |  |  |  | Effect of water, compared to no water |  |  |
| --- | --- | --- | --- | --- | --- | --- | --- |
|  | estimate | exp | s.e. | p | estimate | exp | p |
| water supplementation | -2.330 | 0.097 | 0.363 | <0.001 |  |  |  |
| 0.5% yeast | 0.763 | 2.144 | 0.356 | 0.032 |  |  |  |
| 2% yeast | -1.520 | 0.219 | 0.383 | <0.001 |  |  |  |
| 5% yeast | -2.638 | 0.072 | 0.351 | <0.001 |  |  |  |
| 14% yeast | 2.393 | 10.950 | 0.339 | <0.001 |  |  |  |
| 0.5% yeast * water | 2.358 | 10.571 | 0.545 | <0.001 | 0.028 | 1.029 | 0.958 |
| 2% yeast * water | 2.016 | 7.509 | 0.482 | <0.001 | -0.314 | 0.731 | 0.515 |
| 5% yeast * water | 2.213 | 9.145 | 0.597 | <0.001 | -0.116 | 0.890 | 0.845 |
| 14% yeast * water | -1.564 | 0.209 | 0.549 | 0.004 | -3.894 | 0.020 | <0.001 |

**Table S6.** Effect of diet and water supplementation on mortality within DGRP-853.

| coefficient | Estimates from individual model |  |  |  | Effect of water, compared to no water |  |  |
| --- | --- | --- | --- | --- | --- | --- | --- |
|  | estimate | exp | s.e. | p | estimate | exp | p |
| water supplementation | -0.473 | 0.623 | 0.259 | 0.067 |  |  |  |
| 0.5% yeast | 1.987 | 7.293 | 0.239 | <0.001 |  |  |  |
| 2% yeast | 0.150 | 1.162 | 0.254 | 0.554 |  |  |  |
| 5% yeast | -0.967 | 0.380 | 0.248 | <0.001 |  |  |  |
| 14% yeast | 2.955 | 19.204 | 0.223 | <0.001 |  |  |  |
| 0.5% yeast * water | 0.436 | 1.547 | 0.399 | 0.274 | -0.037 | 0.964 | 0.926 |
| 2% yeast * water | 0.379 | 1.461 | 0.365 | 0.299 | -0.094 | 0.910 | 0.797 |
| 5% yeast * water | 0.680 | 1.973 | 0.373 | 0.068 | 0.206 | 1.229 | 0.58 |
| 14% yeast * water | -2.083 | 0.125 | 0.339 | <0.001 | -2.556 | 0.078 | <0.001 |

**Table S7.** Effect of diet on mortality across 5 water-supplemented DGRP lines (DGRP-195 is reference).

| coefficient | Full Model |  |  |  |
| --- | --- | --- | --- | --- |
|  | estimate | exp | s.e. | p |
| 0.5% yeast | -0.345 | 0.709 | 0.228 | 0.131 |
| 2% yeast | -1.921 | 0.146 | 0.202 | <0.001 |
| 5% yeast | -1.060 | 0.347 | 0.224 | <0.001 |
| 14% yeast | 0.930 | 2.535 | 0.225 | <0.001 |
| 217 | 0.107 | 1.113 | 0.228 | 0.638 |
| 239 | 0.683 | 1.980 | 0.228 | 0.003 |
| 362 | -1.991 | 0.137 | 0.218 | <0.001 |
| 853 | -1.242 | 0.289 | 0.221 | <0.001 |
| 217 * 0.5% yeast | 0.658 | 1.931 | 0.320 | 0.04 |
| 217 * 2% yeast | 0.567 | 1.764 | 0.311 | 0.068 |
| 217 * 5% yeast | 0.096 | 1.101 | 0.321 | 0.764 |
| 217 * 14% yeast | 0.206 | 1.229 | 0.324 | 0.524 |
| 239 * 0.5% yeast | 1.771 | 5.877 | 0.318 | <0.001 |
| 239 * 2% yeast | 2.119 | 8.326 | 0.305 | <0.001 |
| 239 * 5% yeast | 1.042 | 2.834 | 0.321 | 0.001 |
| 239 * 14% yeast | -0.799 | 0.450 | 0.322 | 0.013 |
| 362 * 0.5% yeast | 3.937 | 51.246 | 0.315 | <0.001 |
| 362 * 2% yeast | 2.434 | 11.404 | 0.307 | <0.001 |
| 362 * 5% yeast | 0.691 | 1.995 | 0.318 | 0.03 |
| 362 * 14% yeast | -0.071 | 0.932 | 0.323 | 0.826 |
| 853 * 0.5% yeast | 3.228 | 25.234 | 0.312 | <0.001 |
| 853 * 2% yeast | 2.482 | 11.960 | 0.303 | <0.001 |
| 853 * 5% yeast | 0.739 | 2.094 | 0.322 | 0.022 |
| 853 * 14% yeast | 0.033 | 1.033 | 0.319 | 0.918 |

**Table S8.** Effect of diet on mortality within water-supplemented DGRP-195.

| coefficient | Estimates from individual model |  |  |  |
| --- | --- | --- | --- | --- |
|  | estimate | exp | s.e. | p |
| 0.5% yeast | -0.512 | 0.599 | 0.098 | <0.001 |
| 2% yeast | -2.124 | 0.120 | 0.100 | <0.001 |
| 5% yeast | -1.322 | 0.267 | 0.102 | <0.001 |
| 14% yeast | 1.410 | 4.095 | 0.100 | <0.001 |

**Table S9.** Effect of diet on mortality within water-supplemented DGRP-217.

| coefficient | Estimates from individual model |  |  |  |
| --- | --- | --- | --- | --- |
|  | estimate | exp | s.e. | p |
| 0.5% yeast | 0.357 | 1.429 | 0.214 | 0.095 |
| 2% yeast | -1.487 | 0.226 | 0.215 | <0.001 |
| 5% yeast | -1.011 | 0.364 | 0.216 | <0.001 |
| 14% yeast | 1.260 | 3.527 | 0.215 | <0.001 |

**Table S10.** Effect of diet on mortality within water-supplemented DGRP-239.

| coefficient | Estimates from individual model |  |  |  |
| --- | --- | --- | --- | --- |
|  | estimate | exp | s.e. | p |
| 0.5% yeast | 1.114 | 3.046 | 0.119 | <0.001 |
| 2% yeast | 0.150 | 1.162 | 0.118 | 0.204 |
| 5% yeast | -0.012 | 0.988 | 0.117 | 0.919 |
| 14% yeast | 0.096 | 1.101 | 0.118 | 0.413 |

**Table S11.** Effect of diet on mortality within water-supplemented DGRP-362.

| coefficient | Estimates from individual model |  |  |  |
| --- | --- | --- | --- | --- |
|  | estimate | exp | s.e. | p |
| 0.5% yeast | 3.631 | 37.764 | 0.386 | <0.001 |
| 2% yeast | 0.518 | 1.678 | 0.379 | 0.172 |
| 5% yeast | -0.487 | 0.614 | 0.380 | 0.2 |
| 14% yeast | 0.920 | 2.509 | 0.379 | 0.015 |

**Table S12.** Effect of diet on mortality within water-supplemented DGRP-853.

| coefficient | Estimates from individual model |  |  |  |
| --- | --- | --- | --- | --- |
|  | estimate | exp | s.e. | p |
| 0.5% yeast | 2.811 | 16.623 | 0.222 | <0.001 |
| 2% yeast | 0.564 | 1.759 | 0.216 | 0.009 |
| 5% yeast | -0.284 | 0.753 | 0.216 | 0.19 |
| 14% yeast | 0.923 | 2.518 | 0.216 | <0.001 |

**Table S13.** Effect of diet and water supplementation on fecundity across 5 DGRP lines, derived from linear odel estimates oflog-transformed raw fecundity counts (DGRP-195 is reference). Counts generated using QuantiFly software. Counts are relative, but directly comparable.

| coefficient | Full Model |  |  | Compared to 8% |  |
| --- | --- | --- | --- | --- | --- |
|  | estimate | s.e. | p | estimate | p |
| intercept | 2.731 | 0.067 | <0.001 |  |  |
| water | 0.039 | 0.027 | 0.15 |  |  |
| 0.5% yeast | -1.374 | 0.092 | <0.001 |  |  |
| 2% yeast | -0.595 | 0.092 | <0.001 |  |  |
| 5% yeast | -0.367 | 0.092 | <0.001 |  |  |
| 14% yeast | 0.157 | 0.092 | 0.09 | 2.888 | <0.001 |
| 217 | -0.044 | 0.092 | 0.634 |  |  |
| 239 | -0.331 | 0.095 | 0.001 |  |  |
| 362 | -0.294 | 0.092 | 0.002 |  |  |
| 853 | -0.221 | 0.095 | 0.021 |  |  |
| 217 * 0.5% yeast | 0.113 | 0.130 | 0.387 |  |  |
| 217 * 2% yeast | -0.268 | 0.134 | 0.048 |  |  |
| 217 * 5% yeast | -0.164 | 0.132 | 0.216 |  |  |
| 217 * 14% yeast | -0.224 | 0.132 | 0.092 | -0.110 | 0.404 |
| 239 * 0.5% yeast | 0.509 | 0.139 | <0.001 |  |  |
| 239 * 2% yeast | 0.310 | 0.139 | 0.027 |  |  |
| 239 * 5% yeast | -0.119 | 0.134 | 0.373 |  |  |
| 239 * 14% yeast | -0.262 | 0.136 | 0.056 | -0.436 | 0.001 |
| 362 * 0.5% yeast | 0.524 | 0.130 | <0.001 |  |  |
| 362 * 2% yeast | 0.212 | 0.130 | 0.106 |  |  |
| 362 * 5% yeast | 0.180 | 0.130 | 0.168 |  |  |
| 362 * 14% yeast | -0.202 | 0.130 | 0.122 | -0.339 | 0.009 |
| 853 * 0.5% yeast | 0.249 | 0.134 | 0.065 |  |  |
| 853 * 2% yeast | -0.233 | 0.132 | 0.079 |  |  |
| 853 * 5% yeast | -0.137 | 0.132 | 0.3 |  |  |
| 853 * 14% yeast | -0.657 | 0.134 | <0.001 | -0.721 | <0.001 |

**Table S14.** Effect of diet and water supplementation on fecundity across5 DGRP lines, derived from linear model stimates oflog-transformed mortality-adjusted fecundity counts (DGRP-195 is reference). Counts generated using QuantiFly software. Counts are relative, but directly comparable.

| coefficient | Full Model |  |  | Compared to 8% |  |
| --- | --- | --- | --- | --- | --- |
|  | estimate | s.e. | p | estimate | p |
| intercept | 0.776 | 0.067 | <0.001 |  |  |
| water | -0.004 | 0.027 | 0.873 |  |  |
| 0.5% yeast | -1.379 | 0.093 | <0.001 |  |  |
| 2% yeast | -0.602 | 0.093 | <0.001 |  |  |
| 5% yeast | -0.384 | 0.093 | <0.001 |  |  |
| 14% yeast | 0.148 | 0.093 | 0.114 | 0.925 | <0.001 |
| 217 | -0.050 | 0.093 | 0.594 |  |  |
| 239 | -0.350 | 0.096 | <0.001 |  |  |
| 362 | -0.239 | 0.093 | 0.011 |  |  |
| 853 | -0.241 | 0.096 | 0.013 |  |  |
| 217 * 0.5% yeast | 0.101 | 0.132 | 0.444 |  |  |
| 217 * 2% yeast | -0.301 | 0.136 | 0.028 |  |  |
| 217 * 5% yeast | -0.113 | 0.134 | 0.398 |  |  |
| 217 * 14% yeast | -0.165 | 0.134 | 0.218 | -0.067 | 0.617 |
| 239 * 0.5% yeast | 0.542 | 0.141 | <0.001 |  |  |
| 239 * 2% yeast | 0.311 | 0.141 | 0.028 |  |  |
| 239 * 5% yeast | -0.120 | 0.136 | 0.378 |  |  |
| 239 * 14% yeast | -0.237 | 0.138 | 0.087 | -0.439 | 0.001 |
| 362 * 0.5% yeast | 0.485 | 0.132 | <0.001 |  |  |
| 362 * 2% yeast | 0.177 | 0.132 | 0.182 |  |  |
| 362 * 5% yeast | 0.186 | 0.132 | 0.16 |  |  |
| 362 * 14% yeast | 0.166 | 0.132 | 0.209 | 0.075 | 0.568 |
| 853 * 0.5% yeast | 0.284 | 0.136 | 0.038 |  |  |
| 853 * 2% yeast | -0.227 | 0.134 | 0.091 |  |  |
| 853 * 5% yeast | -0.111 | 0.134 | 0.406 |  |  |
| 853 * 14% yeast | -0.550 | 0.136 | <0.001 | -0.643 | <0.001 |

